# Oxytocin enhances the triangular association among behavioral performance, resting state, and task-state functional connectivity

**DOI:** 10.1101/2022.12.23.521846

**Authors:** Haoming Zhang, Kun Chen, Jin Bao, Haiyan Wu

**Affiliations:** Centre for Cognitive and Brain Sciences and Department of Psychology, University of Macau, Macau, China; Shenzhen Neher Neural Plasticity Laboratory, the Brain Cognition and Brain Disease Institute, Shenzhen Institute of Advanced Technology, Chinese Academy of Sciences (CAS); Shenzhen-Hong Kong Institute of Brain Science-Shenzhen Fundamental Research Institutions, Shenzhen, 518055, China

**Author notes:** Equal contribution (Haoming Zhang, Kun Chen).

## Abstract

The role of oxytocin (OT) in social behavior and social brain networks has been widely documented. However, the effect of OT on the association between social behavior and brain functional connectivity (FC) is yet to be comprehensively explored. In this study, using a face-perception task and multiple connectome-based predictive (CPM) models, we aimed to: 1) determine whether OT could enhance the association between task behavioral performance, resting-state functional connectivity (rsFC), and task-state functional connectivity (tsFC), and 2) if so, determine the role of OT in enhancing this triangular association. We found that both rsFC and tsFC could independently and significantly predict task performance in the OT group, but not in the placebo (PL) group. In addition, the correlation coefficient between rsFC and tsFC was substantially higher in the OT group than in the PL group. The strength of these associations could be partly explained by OT altering the brain’s FCs related to social cognition and face-perception in both resting and task states, mainly in brain regions such as the limbic system, prefrontal cortex (PFC), temporal poles (TP), and temporoparietal junction (TPJ). Together, these results suggest that neuropeptides can increase the consistency of individual differences in different modalities (e.g., behavioral and brain level data).

## 1 Introduction

Oxytocin (OT) is a neuropeptide associated with various social functions [1, 2, 3]. It has been proven to be closely linked to social adaptation and prosocial behaviors [4, 5, 6], and social cognition [7, 8, 9]. For instance, early evidence has shown that OT may increase attention to the eye region of human faces [10] and improve the ability to infer the mental state of others from social cues of the eye region [11]. Further studies observed a more general enhancement effect of OT on motivation or sensitivity to social cues [12, 13, 14], which may manifest in face- or emotion-related tasks. OT can also modulate self- and face-perception, hallmarks of human social cognition [15]. Since OT can increase sensitivity to social stimuli [16, 17], it is reasonable that OT influences facial processing at both the behavioral and neural levels [18, 19].

OT has widely demonstrated its modulation effect in the social brain [20, 21], including the amygdala, anterior cingulate cortex (ACC), prefrontal cortex (PFC), and insula [22, 23]. However, its effects on specific brain areas or functional connectivities are mostly task-dependent and inconsistent across studies [24, 25, 26]. For instance, several studies have shown that OT increases amygdala responses to emotional faces or aversive stimuli [27, 28], while others have observed an attenuation effect in this area [29, 30]. Additional studies have shown that OT changes functional connectivity (FC) in brain regions that belong to the social network [25, 31, 32]. For example, OT increases the effective flow from the midline default network, including the posterior cingulate cortex (PCC) and precuneus, to the salience network, including the ACC and insula, [32] and brain connectivity within the frontal network [33] during the resting-state. OT has also been found to alter brain connectivity strength during different tasks. For instance, OT could enhance the FC between the left amygdala, left anterior insula, and left inferior frontal gyrus in emotional perception and memory tasks [34]. Therefore, the specific modulatory effects of OT on connectivity among social subnetworks have been well-documented.

In addition, OT effects also exhibit individual differences [2, 35], which exist not only at the behavioral level, but also at the brain level [3, 36]. OT could increase correlation between the brain networks in the resting and task states at individual level [37]. However, the specific modulatory effects of OT on the link between individual differences in different modalities (e.g., behavioral and brain levels) remain largely unknown. The connectome-based predictive model (CPM), which predicts a behavior index based on FCs, has been widely used to examine brain-behavior associations [38, 39]. It predicts individual variability in behavior or psychiatric symptoms by extracting and summarizing the most relevant features from FC using full cross-validation [40]. Many prior studies have demonstrated the robustness of CPM [41, 42] in predicting individual differences in fluid intelligence [43], attention [44], creative ability [45], and cheating behavior [46]. Most previous relevant studies used resting state FC to predict behavioral symptoms. In contrast, several recent studies have shown that FC during narrative movie watching [47] or specific tasks [48, 49, 50] provides additional information for CPM prediction.

However, whether and how OT modulates the brain-behavior association is largely unknown. Based on previous progresses on OT effect on behavior and FC, and its individual variability, in the present study, we systematically explored the relationship between social behavior, rsFC, and tsFC, as well as the effect of OT administration on this triangular association. We assessed the following questions: 1) whether the association between behavioral performance and rsFC could be enhanced by OT administration (**Sec. 3.2**); 2) whether the association between behavioral performance and tsFC could be enhanced by OT (**Sec. 3.3**); and 3) whether OT could enhance the similarity between rsFC and tsFC at the whole-brain level (**Sec. 3.4**). For Questions 1-3, we hypothesized that OT could enhance the triangular association between behavioral performance, rsFC, and tsFC. If this hypothesis holds, we then assessed Question 4): how does OT enhance this triangular association (**Sec. 3.5 and Sec. 3.6**)? Since we revealed that OT did not change participants’ behavioral performance in our previous study [51], we then hypothesized that OT enhances the triangular association by altering rsFC or tsFC.

To investigate these questions and test our hypotheses, we first designed a face-perception paradigm with a between-subject design for task fMRI, judging whether the morphed face we present is similar to their own face (see Figure 1 A). In this experiment, we recorded the behavioral performance index, resting state fMRI signals, and task-state fMRI signals. Then, to answer Questions 1 and 2, we used CPM to examine whether FC could predict their corresponding behavioral performance and whether OT could enhance the CPM prediction accuracy of FC on behavioral performance. In this question, CPM was used to determine whether there was a significant association between FC and behavior (Sec. 2.6 CPM predictor). For Question 3, we used a correlation analysis to repeat our previous study [37] at the whole-brain level. For Question 4, we used the CPM to separate the FCs in the OT group from those in the placebo (PL) group and found which FC features contributed more to group differentiation. CPM was used to examine whether there were significant differences between the FCs in the OT and PL groups (Sec. 2.8 CPM classifier). We then examined whether these important features also showed different associations with behavioral performance in the OT and PL groups.

**Figure 1:**
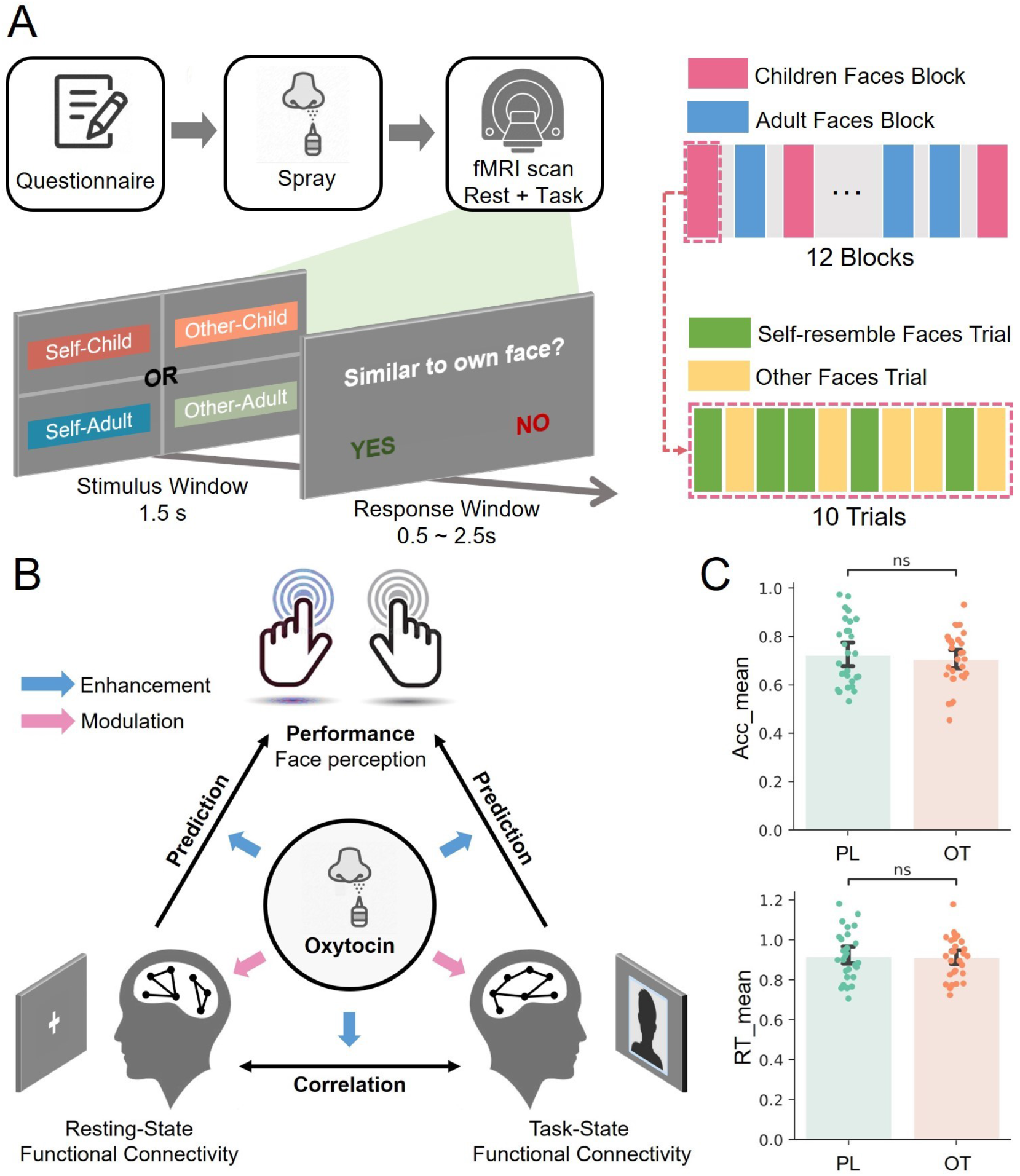
The experimental paradigm, main framework, and behavioral results. (**A**) The experimental paradigm. Participants need to judge whether the morphed face is similar to their own. There are four conditions of faces: Self-Child condition was the face morphed using the participant’s own face and a stranger child’s face; faces of the Other-Child conditions were morphed using an adult stranger’s face and a stranger child’s face; faces in the Self-Adult condition were morphed using the participant’s own face and a stranger adult’s face; faces in the Other-Adult condition were morphed using two adult strangers’ faces. (**B**) The main structure of the present study. (**C**) The behavioral results of the PL and OT groups. From top to bottom, we present the behavioral results of averaged accuracy (Acc mean) and average reaction time (RT mean). There were no significant behavioral differences between the groups.

## 2 Methods

### 2.1 Participants

Fifty-nine healthy male participants (age: mean ± SD = 20.9 ± 2.32 years old) were recruited by online advertisement. All participants were right-handed, with normal or corrected-to-normal vision. All participants signed an informed consent form before the formal experiment. Patients were only included if they were confirmed that they were not suffering from any significant medical or psychiatric illness, not currently using the medication, and were not consuming alcohol or smoking daily. The experimental protocol was approved by the local ethics committee of the Beijing Normal University.

### 2.2 Drug administration

We used a double-blind placebo-controlled group design to investigate the effects of a single dose of intranasal OT (24 IU) [52] on functional connectivity (FC) and corresponding behavioral performance in a face-perception task. The participants (all males) were randomly assigned to the OT group (n = 30) or PL group (n = 29). More details regarding the treatment can be found in our previous studies [37, 33, 51].

### 2.3 Experimental paradigm

The experimental paradigm was a face-perception task with a morphed face, following previous studies [53] (Figure 1 A). Specifically, we used a face-perception task with face stimuli that morphed the photos of an adult or child onto the participant or another stranger [51]. For each trial, the target-morphed face was presented for 1.5 s. Participants were then asked to judge whether the face resembled their own faces in the following response window within 0.5–2.5 s. There were a total of 12 blocks: six were morphed children’s faces blocks and six were morphed adult faces blocks. Each block contained 10 trials, five of which presented a self-resembling face, and five of which presented other-resembling faces. Thus, there were four types of facial stimuli. The self-resembling faces were created by morphing the participant’s face with a face of a 23-year-old adult (self-adult) or a face of a 1.5-year-old child (self-child). The other-resembling faces were created by morphing a stranger’s face with the face of a 23-year-old adult (other-adult) or a 1.5-year-old child (other-child). All facial expressions were neutral.

### 2.4 MRI acquisition

All MRI data were acquired using a 3.0 T Siemens Tim Trio scanner equipped with a 12-channel head coil. First, high-resolution T1-weighted images were acquired for each participant (TR = 1.9 *s*, TE = 2.15 *ms*, flip = 9°, FOV = 256 *mm*, 176 sagittal slices, slice thickness = 1 *mm*). We collected resting state and task-state fMRI data successively. All fMRI data were collected using an echo-planar imaging sequence (TR = 2 *s*, TE = 40 *ms*, flip = 90°, FOV = 210 *mm*; 128 × 128 matrix, 25 contiguous 5 *mm* slices parallel to the hippocampus and interleaved).

### 2.5 fMRI preprocessing

fMRI data preprocessing was performed using SPM12 (Statistical Parametric Mapping; https://www.fil.ion.ucl.ac.uk/spm/software/spm12). The functional image time series were preprocessed to compensate for motion correction, slice-timing correction, and linear detrending; thereafter, they were co-registered to the T1-weighted anatomical image, normalized to Montreal Neurological Institute space, and smoothed with an isotropic Gaussian kernel of 6 mm full width at half maximum. Finally, the fMRI data were high-pass filtered with a cutoff of 0.01 Hz. White matter, cerebrospinal fluid (CSF), global, and six head motion parameters, as well as their squares, derivatives, and squares of derivatives, were regressed [54]. The resulting residuals were then low-pass filtered with a cutoff of 0.1 Hz.

### 2.6 CPM predictor

The main analysis utilized CPM models to predict behavioral indices or participant groups based on rsFC or tsFC. The CPM predictor model built a bridge between FC and behavior for each group. The workflow of the CPM predictor model is shown in Fig. 2. First, we extracted the averaged blood-oxygen-level-dependent (BOLD) time series of the 90 brain regions based on the AAL atlas [55] as ROIs. A 90 × 90 FC matrix was obtained by Pearson’s correlation between the averaged BOLD time series of each pair of ROIs. To remove diagonal and repetitive features, we retained only the lower-triangle matrix (4005 FC features) for further analysis. The Nilearn toolbox (https://nilearn.github.io/) [56] was used to construct the FC matrix.

**Figure 2:**
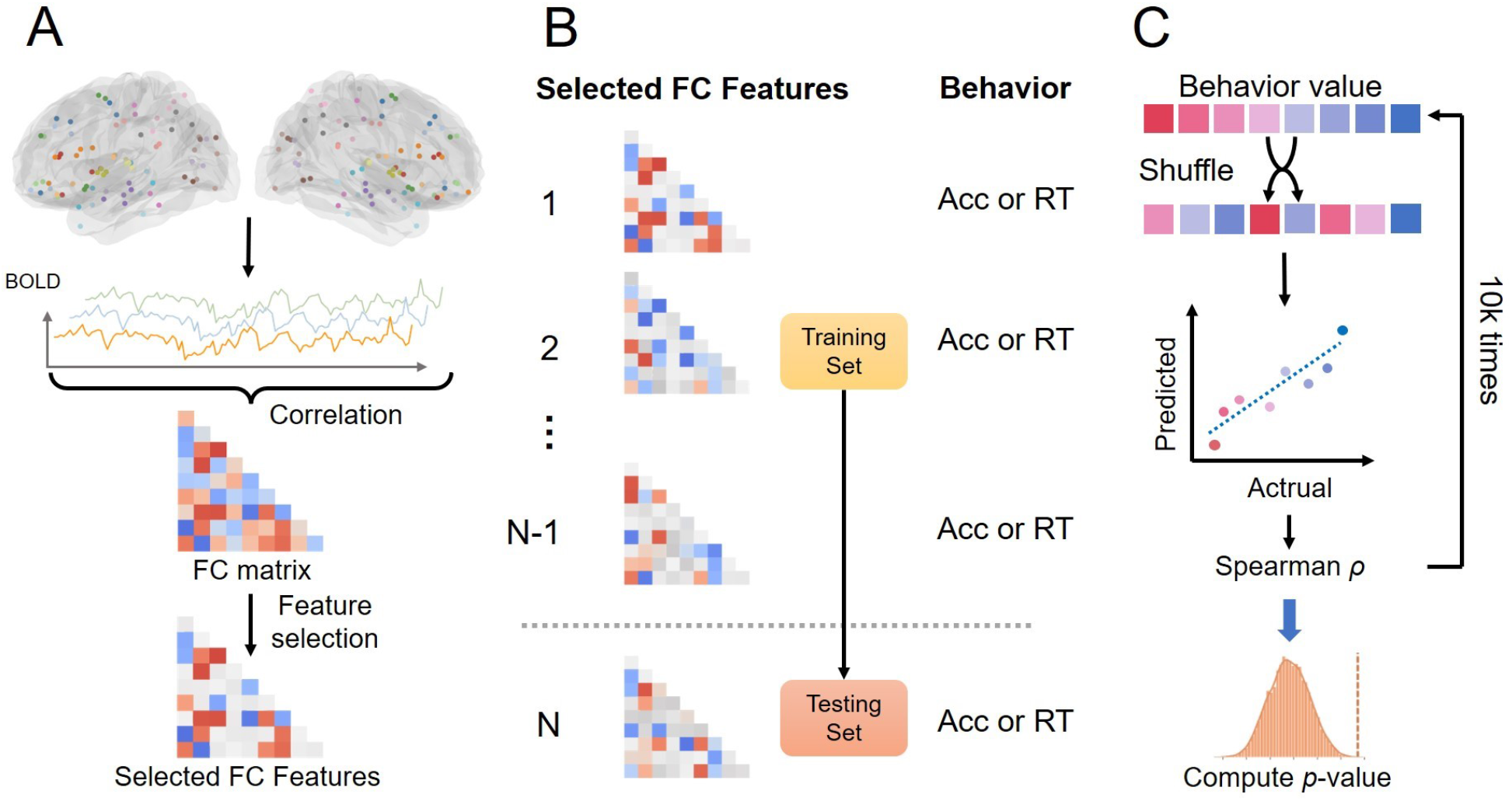
The workflow of the CPM predictor analysis for the present study. (**A**) FC matrix construction and feature selection. (**B**) Model validation with LOOCV. (**C**) Model performance measurement by Spearman correlation and statistics calculated by Permutation test.

Before the actual CPM, we selected the features that had a significant Pearson correlation between FC and the behavioral index (see Figure 2 A), which was used to select the features with potential information for prediction and was consistent with previous studies [40, 46, 45]. Only FCs whose P-value of Pearson’s correlation was lower than the threshold were maintained. By changing the threshold value, different FC quantities can be retained. We controlled the quantity of FC within a reasonable range (10-100), neither too little, resulting in insufficient information, nor too much, resulting in over-fitting of the regression analysis later. Herein, we used two different thresholds: *p* =0.05 and *p* =0.01.

We then implemented a support vector machine (SVM) regressor with a linear kernel to predict participants’ behavioral index based on their FC and validated it with leave-one-out cross-validation (LOOCV) (Figure 2 B) [45, 57, 38]. For each validation, the prediction model was fitted based on *n*−1 (*n* is the number of participants in each group) participants’ selected FCs and their corresponding behavioral index. The model was then tested on the leave-out participants’ data to obtain a predicted behavior value. Thus, LOOCV resulted in *n* behavior index predictions for *n* participants. We used the Spearman’s correlation between the predicted and actual behavioral values as the model accuracy measurement. Additional details of CPM predictor parameters are provided in Table S1.

After obtaining accuracy by LOOCV, we examined whether the model performance was significantly higher than chance. We used a permutation test to evaluate the statistical significance of the CPM prediction. We randomly shuffled the behavioral index values 10,000 times and used LOOCV to obtain the corresponding accuracy for each shuffle (Figure 2 C). Finally, we sorted the accuracy from 10,000 permutations and counted the position of the true model accuracy to obtain the p-value.

### 2.7 FC similarity analysis

Since the similarity between individuals’ rsFC and tsFC could be enhanced by OT has been strictly proven [37], we used a simple method to replicate the effect of OT on the association between rsFC and tsFC. We performed the same correlation analysis for both groups (OT/PL). First, we extracted the lower-triangle FC matrix for each state (resting/task) and group. For each group, we calculated the Pearson correlation coefficient between the resting and task states for each FC edge. We then obtained pairs of FC edge correlation coefficients representing the similarity between rsFCs and tsFCs for each group. Finally, we used pairwise comparison statistics (paired t-test) to investigate whether there was a significant difference in the resting-task FC similarity between the OT and PL groups.

### 2.8 CPM classifier

The CPM classifier was used to classify OT/PL groups based on FC patterns and identify FCs with significant differences between the OT and PL groups. The CPM classifier followed a workflow similar to that of the CPM predictor, but with one classifier across both groups (Figure S2). The same FC extraction approach was used to obtain a lower-triangle FC matrix between the AAL ROIs based on Pearson correlation. Feature selection was slightly different from that in the predictor workflow. For each FC in the lower-triangle matrix, we conducted logistic regressions between each FC value and their group label (OT/PL) to obtain the regression coefficient. Only the FCs whose P-value of the regression coefficient was lower than the threshold were retained. By changing the threshold value, different FC quantities can be retained. In the present study, we used three different thresholds, namely *p* =0.05, *p* =0.02, and *p* =0.01.

Then, we implemented a linear kernel SVM classifier based on the maintained FCs to classify the corresponding groups (OT/PL) using LOOCV [45, 57, 38]. More specifically, the model was fitted based on *n*−1 (*n* is the total number of participants in the two groups) participants’ FC matrixes and corresponding group labels. The fitted model was tested on one left-out participant. After *n* cycles, each participant was tested once and eventually achieved accuracy, which was used to quantify the performance of the model. Finally, we used the same permutation procedure but shuffled the group labels rather than the behavior indices.

To quantify the contribution of each FC in the CPM classifier model, we developed a similar lesion-based approach [46, 58] to explore which FC contributes the most to classification accuracy. Specifically, we removed each FC and retained the remaining FCs. We used the remaining FCs to train and test the model using the same LOOCV process, and obtained the corresponding accuracy of the model that lost this FC. Feature importance was then calculated as the difference in accuracy between the full model (no FC’s removed) and the model without the corresponding FC. Consequently, the higher the difference, the more important the FC.

### 2.9 Regression analysis

After finding the important FCs in the classification, we speculated whether these FCs may only be significantly associated with behavioral performance in the OT group. To achieve this goal, we performed linear regression analysis on the OT group and the PL group, respectively. The behavioral performance index was linearly regressed onto functional connectivity (Eq. 1).

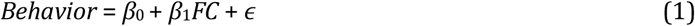

Where *FC* denotes the selected and important functional connectivity. *β*_0_ as the regression intercept. *β*_1_ denotes the regression slope. *Behavior* denotes behavioral performance value. The model parameters were estimated using ordinary least squares. After each regression process, we obtained the regression slope coefficient and corresponding p-value coefficient.

The corresponding codes for all analyses in the present study will be available online at http://github.com/andlab-um/OT-cpm.

## 3 Results

### 3.1 Behavioral results

We used a face-perception task in our study. The participants were required to judge whether the morphed face was similar to their own. In the face-perception task, we calculated two indices for each participant: mean accuracy (Acc mean) and average RT mean. Figure 1 C presents the two behavioral response indices from the OT and PL groups in the face-perception task. We performed pairwise comparison statistics (t-test) between the OT and PL groups for these behavioral indices and found no significant difference between the two groups.

### 3.2 OT enhances the prediction effect from rsFC to behavior

First, we assessed whether OT-enhanced the association between rsFC and behavior. We used a CPM predictor model to predict the behavioral index based on rsFCs for the OT and PL groups and examined whether the prediction performance of the CPM predictor in the OT and PL groups differed. We trained two models with the same hyperparameters for each behavioral index to compare the predictive abilities of the two groups. Because there were two groups (OT/PL) and two behavioral performance indices (Acc mean/RT mean), we trained four CPM predictor models in total.

The first part of the CPM predictor uses the Pearson correlation to select significant features from all FC features. To predict the Acc mean, we selected 57 features in the OT group and 18 features in the PL group by setting the threshold to 0.01. To predict the mean RT, 35 features in the OT group and 80 features in the PL group were selected by setting the threshold to 0.05 (Table S1).

Based on the selected FCs, we used a linear SVM regressor to predict task performance. We also used LOOCV and Spearman’s correlation to obtain predictive performance and used permutation for significance testing. The results for the CPM predictor are plotted in Figure 3. We found that a significant predictive effect existed only in the OT group. Figure 3 A presents the results of using the CPM model to predict the ACC mean. The prediction effect was insignificant in the PL group (*ρ* = 0.101, *p* = 0.347). However, in the OT group, we observed significant prediction accuracy (*ρ* = 0.492, *p* = 0.0057**). Figure 3 B presents the results of using the CPM model to predict mean RT; the prediction effect was not significant in the PL group (*ρ* = −0.411, *p* = 0.943) but significant in the OT group (*ρ* = 0.677, *p <* 0.001***).

**Figure 3:**
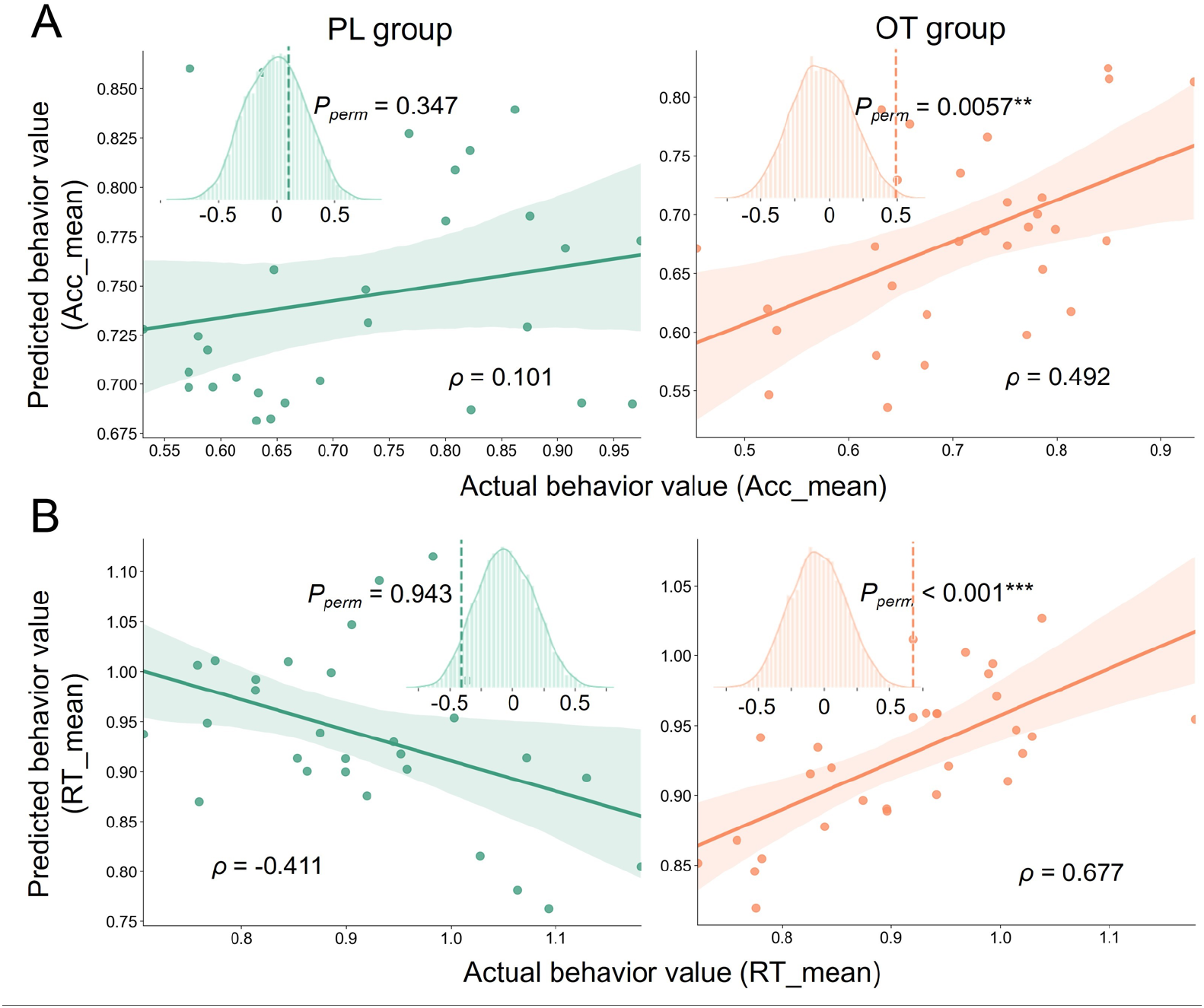
Results of using the CPM predictor to predict task performance by resting state FC. We present the Spearman correlation between the actual behavioral value and predicted behavioral value alongside the permutation result of the correlation coefficient. (A) The result of using the CPM predictor to predict Acc mean in the PL (left) and OT (right) groups. (B) The result of using the CPM predictor to predict RT mean in the PL (left) and OT (right) groups.

### 3.3 OT enhances the prediction effect from tsFC to behavior

Next, we tested whether there was a similar pattern between tsFC and behavior. We used the same CPM prediction approach to predict the behavior index of each group based on the tsFCs. The first part is feature selection. To predict the Acc mean, we selected nine features in the OT group and 146 features in the PL group by setting the threshold to 0.05. To predict the mean RT, 314 features in the OT group and 68 features in the PL group were selected by setting the threshold to 0.01 (Table S1).

Figure 4 A and B presents the results of using the CPM model to predict ACC mean and RT mean, respectively. Similar to the previous result of rsFC, a significant prediction effect only existed in the OT group. The prediction effect from tsFC to ACC mean was not significant in the PL group (*ρ* = −0.432, *p* = 0.347) (Figure 4 A). Furthermore, there is a positive relationship between the actual behavioral value and prediction in the OT group, although the effect showed only marginal significance (*ρ* = 0.366, *p* = 0.052). A pattern similar to the rsFC CPM was also found in the prediction from task FC to RT mean (Figure 4 B). The prediction effect was not significant in the PL group (*ρ* = −0.049, *p* = 0.331) but significant in the OT group (*ρ* = 0.335, *p* = 0.024*). Results demonstrated that OT-enhanced the prediction effect from FCs to behavior indices in both the resting and task-state fMRI.

**Figure 4:**
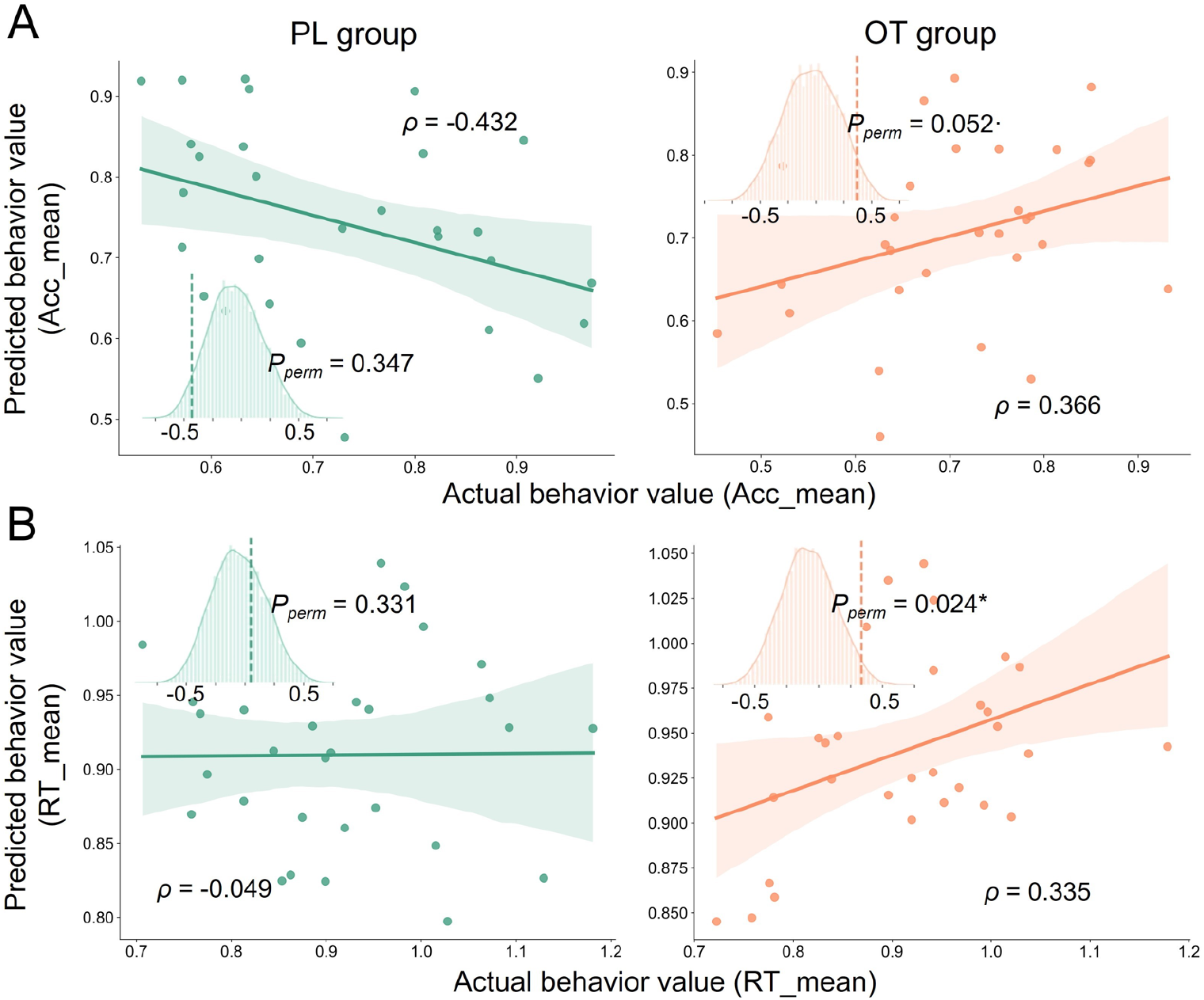
Results of using CPM predictor to predict task performance by task-state FC. The Spearman correlation between actual behavioral value and predicted behavioral value alongside the permutation result of the correlation coefficient. (A) The result of using the CPM predictor to predict Acc mean in the PL (left) and OT (right) groups. (B) The result of using the CPM predictor to predict RT mean in the PL (left) and OT (right) groups.

### 3.4 OT enhances similarity between rsFC and tsFC

Our previous research has proven that the similarity between individuals’ rsFC and tsFC could be enhanced by OT [37]. In the present study, we used a simple method to verify that OT repeatedly enhances resttask association at the whole-brain level. Figure S3 shows that the correlation coefficient between rsFC and tsFC in the OT group was significantly higher than that in the PL group (*p <* 0.001), which is consistent with our previous work.

### 3.5 OT alters resting state functional connectivity

After proving that OT could enhance the association between behavioral performance, rsFC, and tsFC, we were curious about how OT enhances this triangular association. First, OT may alter one or more vertices of the triangle. Since behavioral performance was not significantly different between the OT and PL groups (Figure 1 C) we hypothesized that OT may directly alter the participants’ FC. To prove this conjecture, we developed the CPM classifier to test whether it could discriminate the FCs in the OT group from the PL group and determine which FC features were significantly different between the two groups. The first part uses logic regression to select significant features from a 90 × 90 FC matrix. After this process, 25 significant FCs values were retained by setting the threshold value to *p <*0.02. To verify the stability of the results, we also used two different thresholds to extract the FC features and conducted the following analysis separately (Supplementary Section 1; Figure S4). We used an SVM classifier based on the selected FCs to classify the groups (OT/PL). We also used LOOCV to obtain accuracy and a permutation test to obtain p-values. We found that the classification accuracy was significantly higher than the chance level: the classification accuracy resulted in *Acc* = 0.678 with a significance level of *p* = 0.0047** (Figure 5 A).

**Figure 5:**
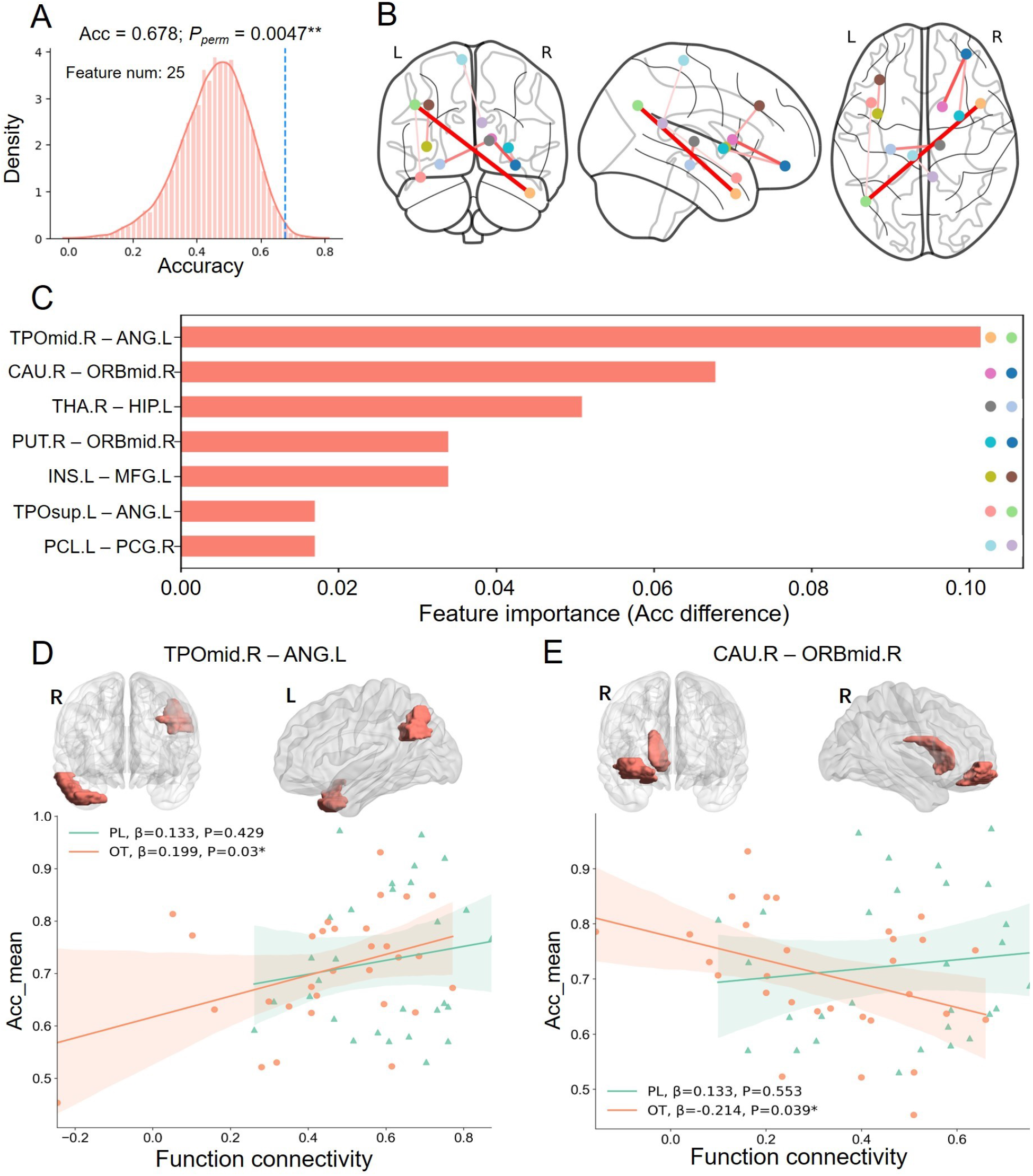
Result of the CPM classifier in resting state FC classification. (A) The permutation result of the model performance. (B) The location of the top seven important FC features. (C) The feature importance of the top seven FC features. (D) Association between Acc mean and TPOmid.R ANG.L. The brain map above shows the location of the two ROIs of FC, and the correlogram below shows the association between FC and Acc mean in the PL (Green line) and OT (Orange line) groups. (E) Association between Acc mean and FC Cau.R ORBmid.R. The brain map above shows the location of the two ROIs of FC, and the correlogram below shows the association between FC and Acc mean in the PL (Green line) and OT (Orange line) groups.

Next, we explored which FCs contribute the most to classification accuracy and found that seven FC features positively impact classification performance. We revealed that the FC between the right temporal pole middle temporal gyrus (TPOmid.R) and the left angular gyrus (ANG.L) is crucial for classifying whether FC belongs to participants treated by OT or PL. In addition, connectivity between the right caudate nucleus (CAU.R) and the right middle frontal gyrus orbital part (ORBmid.R), the right thalamus (THA.R) and the left hippocampus (HIP.L), the right lenticular nucleus putamen (PUT.R), the right middle frontal gyrus orbital part (ORBmid.R), the left insula (INS.L) and the left middle frontal gyrus (MFG.L), the left temporal pole superior temporal gyrus (TPOsup.L), the left angular gyrus (ANG.L), the right and left paracentral lobule (PCL.L), and the posterior cingulate gyrus (PCG.R) were found to be important FCs (Figure 5 C). The locations of all important FCs are shown in Figure 5 B.

Since we proved that OT affects rsFC and found significantly different FCs between the OT and PL groups, we speculated whether these rsFCs that OT significantly alters will show different associations with behavioral performance. For each pair of rsFC and behavior, we performed a regression analysis using rsFC as the regressor and task behavior performance as the data to be regressed. Regression analysis was performed separately for the OT and PL groups to obtain the respective regression coefficients in the two groups. We only focused on rsFC, which showed significant differences between OT and PL, so we chose the first three rsFC features that contributed most to the CPM model (TPOmid.R - ANG.L; CAU.R - ORBmid.R; and THA.R - HIP.L).

The PL group found no significant association between rsFC features and task behavior. However, in the OT group, we found that the first two rsFCs were associated with the behavioral index. The FC between the right temporal pole of the middle temporal gyrus (TPOmid.R) and the left angular gyrus (ANG.L) was significantly associated with the mean Acc (*β* = 0.199, *p* = 0.03*) in the OT group, but not in the PL group (*β* = 0.133, *p* = 0.429). The FC between the right caudate nucleus (CAU.R) and the orbital part of the right middle frontal gyrus (ORBmid.R) was significantly negatively associated with the mean Acc (*β* = −0.214; *p* = 0.039*) in the OT group, but not in the PL group (*β* = 0.133; *p* = 0.553). We further used a statistical model to examine whether OT had a moderating effect on the association between rsFC and behavior [59] and found a marginally significant moderating effect (*p* = 0.081; Supplementary Section 3).

### 3.6 OT alters task-state functional connectivity

Next, we performed the same CPM classification analysis for task-state FC and obtained a classification accuracy significantly higher than chance (*Acc* = 0.661; *P* = 0.0381*; Figure 6 A). Similar to the classification of rsFC, we used two different thresholds to perform the same analysis (Supplementary Section 2; Figure S5). The results indicated that OT-altered FC in both resting and task states. Similarly, we identified four positively influential FC’s that contributed the most to task-state OT/PL classification (Figure 6 B). We showed FC between the right temporal pole superior temporal gyrus (TPOsup.R) and left superior frontal gyrus orbital part (ORBsup.L), TPOsup.R, and the left supramarginal gyrus (SMG.L), and the left temporal pole middle temporal gyrus.

**Figure 6:**
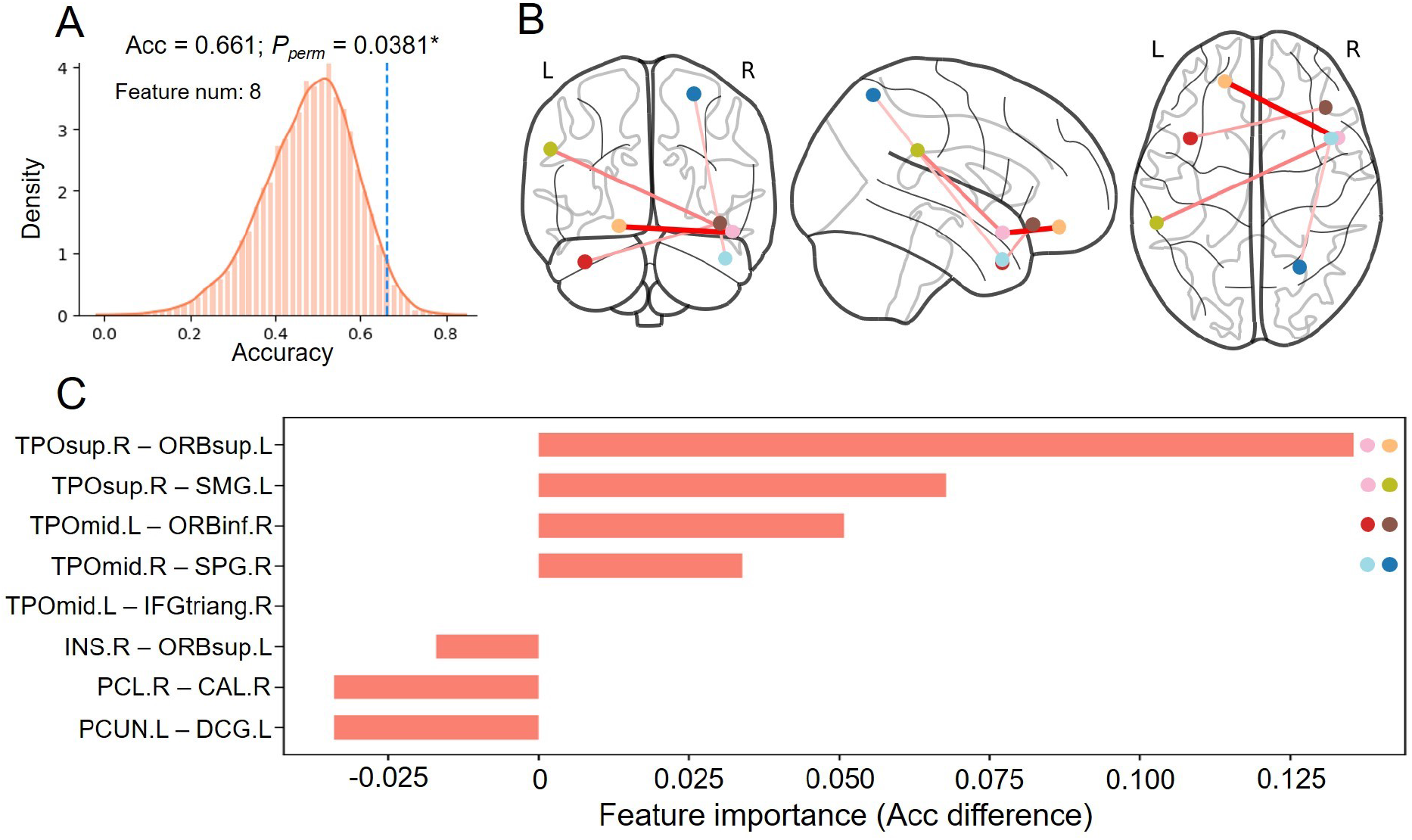
Result of the CPM classifier in task-state FC classification. (A) The permutation result of the model performance. (B) The location of the top four important FC features. (C) The feature importance of all eight FC features.

(TPOmid.L) and inferior frontal gyrus orbital part (ORBinf.R), the right temporal pole middle temporal gyrus (TPOmid.R), and the right superior parietal gyrus (SPG.R) exhibited a significant difference between the OT and PL groups in the task-state (Figure 6 C). For rsFC, we found that only one of the four tsFCs showed a different association with behavioral performance between the OT and PL groups. The FC between the right temporal pole superior temporal gyrus (TPOsup.R) and the left supramarginal gyrus (SMG.L) was significantly associated with the mean RT (*β* = 0.204; *p* = 0.04*) in the OT group, but not in the PL group (*β* = −0.127; *p* = 0.414) (Figure S6). The modulating effect was marginally significant (*p* = 0.064; Supplementary Section 3).

## 4 Discussion

In the present study, we conducted a series of whole-brain level analyses to investigate the modulatory effects of OT on behavioral performance, rsFC, and tsFC. Using CPM-based methods, our findings demonstrate, for the first time, that OT could enhance the association between behavioral performance and FC in both resting- and task-state. OT was also found to enhance the similarity between rsFC and tsFC. Furthermore, our results showed that OT enhances this triangular association (Figure 1 B) by altering the FC in social-related areas in both the resting and task states. Overall, our work validated the triangular association enhancement effect of OT and provided the first evidence that OT could enhance the association between FCs and behavior performance by altering FCs in both resting and task states.

Utilizing CPM predictor models, we demonstrated that FCs in both resting and task states could significantly predict Acc mean and RT mean in the OT group but not in the PL group (Questions 1 and 2). This result indicates that OT can enhance the brain-behavior association. Brain-behavior association is generally considered the association between individual differences in the brain activity and behavior [60]. Several studies have confirmed that OT modulates brain activity depending on individual characteristics, such as age [61], gender [62], and oxytocin receptor (OXTR) genetics [63]. Therefore, OT may increase individual differences in social brain activity related to self-referential or face recognition, which resulted in a stronger brain-behavior association in our task. Our analysis also revealed that OT significantly increased the similarity between rsFC and tsFC (Question 3), which is in line with previous studies showing robust coupling between resting state measures and task-induced brain activity [64, 65]. In addition to enhancing the coupling between rsFC and tsFC, rsFC showed better prediction precision than tsFC for both behavioral indices in the OT group. This may indicate that face perception accuracy and reaction time represent some inherent attributes of individuals, which can be more easily captured by rsFC [66].

Further analysis supported our hypothesis that OT might alter FC to enhance the triangular association (Question 4). The results of the CPM classifier demonstrated that OT could significantly alter FC in both resting and task states. It also revealed that OT has a more significant influence on rsFC than tsFC because the CPM classifier of the rsFC shows higher classification accuracy, stronger significance, and more stable classification results (Supplementary material). Since a previous study found that rsFC contains more stable traits of individuals [66], our results further indicate that OT mainly causes changes in the individuals’ baseline activity instead of task-related activity[25]. Notably, our results showed a consistent relationship between the performance of behavior prediction and group classification performance. More specifically, compared to tsFC, OT had a more significant impact on rsFC, which could better predict behavioral performance after OT administration. This result may also prove that the enhancement of triangular association is related to the alteration of FCs.

OT could alter FCs belonging to the social network, which has been well-documented in previous studies [25, 31, 32]. For instance, previous studies have found that OT-altered FCs between the limbic system (e.g., amygdala, ACC, hippocampus), orbitofrontal cortex, precuneus, SMG, and middle temporal sulcus [67]. Another study also found that brain connectivity in regions comprising the thalamus and superior temporal sulcus might be modulated by OT [68]. Relatedly, for rsFC, we demonstrated that the FC of brain regions in TP (TPOmid.R; TPOsup.L), PFC (ORBmid.R; MFG.L), TPJ (ANG.L), insula (INS.L), and limbic system (CAU.R; THA.R; HIP.L) made significant contributions to the classification of OT and PL. For tsFC, we demonstrated that the FC of brain regions in the TP (TPOmid; TPOsup.R), PFC (ORBsup.L; ORBinf.R), and TPJ (SMG.L; SPG.R) is important for classification. Our findings are consistent with those of previous studies [67, 68] and further provide evidence, at the whole-brain level, that the FC between these social brain regions could be regulated by OT [25].

Moreover, our regression analysis demonstrated that OT-enhanced the brain-behavior association by altering FC. Several important FCs in the classification were only significantly associated with behavioral performance in the OT group, similar to the result in the CPM predictor. For the resting-state, the FC between the TP (TPOmid.R) and TPJ (ANG.L) and the FC between the caudate (CAU.R) and vmPFC (ORBmid.R) was significantly associated with the mean Acc after OT administration. Previous studies have found that the TPJ and vmPFC are critical regions in social cognition [69] and have recognized the role of TP in face recognition and social information processing [70]. In addition, lesions in the connection between the caudate and vmPFC result in impaired social cognition [71, 72]. Therefore, these FCs may be associated with social perception and behavioral performance. For the task-state, the FC between the TP (TPOsup.R) and TPJ (SMG.L) was significantly associated with the mean RT after OT administration. SMG was found to be related to awareness, which helps control attention and reduce reaction time [73]. Notably, not all important FCs in classification show differences in brain-behavior associations between the OT and PL groups, indicating no complete causal relationship between OT-altered brain FC and OT-enhanced brain-behavior association. Notably, not all important FCs in classification show differences in brain-behavior associations between the OT and PL groups, indicating no complete causal relationship between OT-altered brain FC and OT-enhanced brain-behavior association. Besides, the brain-behavior association may reflect mapping from the unique FC pattern instead of every single FC to the behavioral performance, so only part of FCs showed significance in regression analysis.

Our research demonstrates that OT affects the similarity between different modalities rather than a single modality (only behavior or brain activity). Although many studies have explored the consistency between individual differences in different modalities [38, 45, 49], including our previous studies [74, 75, 76], few studies have investigated the influence of neuropeptides, including OT, on this consistency. Mental diseases are also closely related to social abilities and individual differences [77]. OT has been tested as a treatment option for psychopathologies related to social dysfunctions [78], such as autism spectrum disorders, anxiety disorders, social phobia, and schizophrenia [79]. However, the treatment effect of OT shows substantial individual variation and may depend on many other factors [80]. To address this controversy, our study may provide a new understanding of the individual differences caused by OT. If OT enhances the association between different modalities, it is important to pay attention to other therapies, such as behavioral or brain stimulation therapy, while applying OT treatment.

Our study had limitations and there are several potential avenues for future research. First, our current sample was limited to male participants, and we did not test the results by implementing other behavioral experimental tasks. Second, previous studies showed that tsFC might be more suitable for dynamic FC calculation than the current static FC calculation, considering the high time-varying property of the task-state BOLD signal [81]. Dynamic methods have been demonstrated to better characterize BOLD signals at a higher temporal resolution [82, 83], which can explain more behavioral variance than static FC in a global manner [84, 85] and may result in higher prediction accuracy. Third, the current results are more similar to the OT effect on brain-behavioral prediction in the short term, since resting fMRI scanning immediately follows the task. Future work may refine our findings by investigating other social tasks with short- and long-term predictions in OT and PL groups. Because of the constraints of fMRI research, future work should combine fMRI with other approaches, such as EEG and MEG, to build a high temporal resolution association between behavioral and neural dynamics. Finally, although it helps reveal many aspects of brain-human behavior association, CPM is a data-driven approach in neuroimaging that can predict rather than explain. Future theory-driven work can provide insights into the association between increased brain-behavior and OT treatment.

## 5 Conclusion

In conclusion, our results provide the novel insight that OT may enhance the triangular association between face-perception behavior, rsFC, and tsFC. This enhancement could be partly explained by OT altering the FC between social brain regions in both resting and task states. This result provides evidence that neuropeptides can enhance consistency between individual differences in different modalities. Future studies should be conducted to validate this effect of OT using different social tasks, neuroimaging signals, or patients with mental disorders.

## Acknowledgement

This work was mainly supported by the Science and Technology Development Fund (FDCT) of Macau [0127/2020/A3, 0041/2022/A], the Natural Science Foundation of Guangdong Province (2021A151501 2509), Shenzhen-Hong Kong-Macao Science and Technology Innovation Project (Category C) (SGDX2020110309280100), MYRG of University of Macau (MYRG2022-00188-ICI) and the SRG of University of Macau (SRG202000027-ICI). We also thank all research assistants who provided general support in participants recruiting, and data collection.

## Supplementary Materials

### 1 CPM classify result of rsFC with other p-value threshold in feature selection

The CPM classifier’s first part uses logic regression to select significant features from the 90*90 FC matrix. To verify the stability of the results, we used another two different thresholds (*p <*0.05, *p <*0.02) to extract FC features and conduct the next analysis separately. When we set the P-value threshold of regression as *p <*0.05, 131 significant FCs could be retained. When we set the threshold value as *p <* 0.01, there are still 5 significant FCs could be maintained.

We respectively perform the above analysis on two group of FCs selected under two different P-value thresholds, and found they both showed significant classification accuracy: For 5 FCs selected by *p <*0.01, the classification accuracy resulted in *Acc* = 0.627 with significant level *p* = 0.0371* (see Figure S4 A). For 131 FCs selected by *p <*0.05, the classification accuracy resulted in *Acc* = 0.661 with significant level *p* =0.0081** (see Figure S4 B).Based on the highest accuracy and proper number of FC features, we used the 25 FCs selected by *p <*0.02 in the following analysis.

### 2 CPM classify result of tsFC with other p-value threshold in feature selection

Same to the classification analysis in rsFC, we used another two different thresholds (*p <*0.05, *p <*0.02) to extract FC features and conduct the next analysis separately. When we set the P-value threshold of regression as *p <*0.05, 42 significant FCs could be retained. When we set the threshold value as *p <* 0.01, there are no significant FCs could be maintained.

We respectively perform the above analysis on the 42 FCs selected by *p <*0.02, and found it showed significant classification accuracy: For 5 FCs selected by *p <*0.01, the classification accuracy resulted in *Acc* = 0.627 with significant level *p* = 0.0371* (see Figure S4 A).

### 3 Moderating effect test

We further examined the moderating effect of OT through statistical models (Figure S2 D). More specifically, whether the relation between FC and behavioral performance would be significantly affected by OT. We use multiple regression analysis. We included Group (OT/PL) as a moderator variable, the product of FC and Group (FC * Group) as the interaction term. If the slope coefficient of the interaction term is significant, it indicates that there is a significant moderating effect. The regression model used to test the moderating effect is as follows:

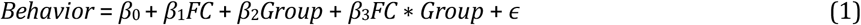

Where *β*_0_ is the regression intercept. *β*_1_ as the regression slope of FC. *β*_2_ as the regression slope of Group (OT/PL). *β*_3_ as the regression slope of interaction effect variable FC * Group and provides an estimate of the moderating effect. If *β*_3_ is statistically different from zero, it would prove that OT has a significant moderating effect.

For the association between Acc mean and rsFC of TPOmid.R - ANG.L, the moderating effect of OT is insignificant (*p* = 0.713). For the association between Acc mean and rsFC of CAU.R - ORBmid.R, the moderating effect of OT is marginal significant (*p* = 0.081·). For the association between Acc mean and tsFC of TPOsup.R – SMG.L, the moderating effect of OT is insignificant (*p* = 0.064·)

**Table S1:** Details of CPM predictor parameters. p-value: P-value threshold for feature selection; OT F: features for OT group; PL F: features for PL group

| Behavior data | FC data | p-value | OT F | PL F | Kernal | Other SVM parameter |
| --- | --- | --- | --- | --- | --- | --- |
| ACC mean | Rest | 0.01 | 57 | 18 | Linear | Default |
| RT mean | Rest | 0.05 | 35 | 80 | Linear | Default |
| ACC mean | Task | 0.05 | 9 | 146 | Linear | Default |
| RT mean | Task | 0.01 | 314 | 68 | Linear | Default |

**Table S2:** Details of CPM classify parameters. p-value: P-value threshold for feature selection

| FC data | p-value | Features | Kernal | Other SVM parameter |
| --- | --- | --- | --- | --- |
| Rest | 0.01 | 5 | Linear | Default |
| Rest | 0.02 | 25 | Linear | Default |
| Rest | 0.05 | 131 | Linear | Default |
| Task | 0.01 | *none* | Linear | Default |
| Task | 0.02 | 8 | Linear | Default |
| Task | 0.05 | 42 | Linear | Default |

## 4 Tables and Figures

**Figure S1:**
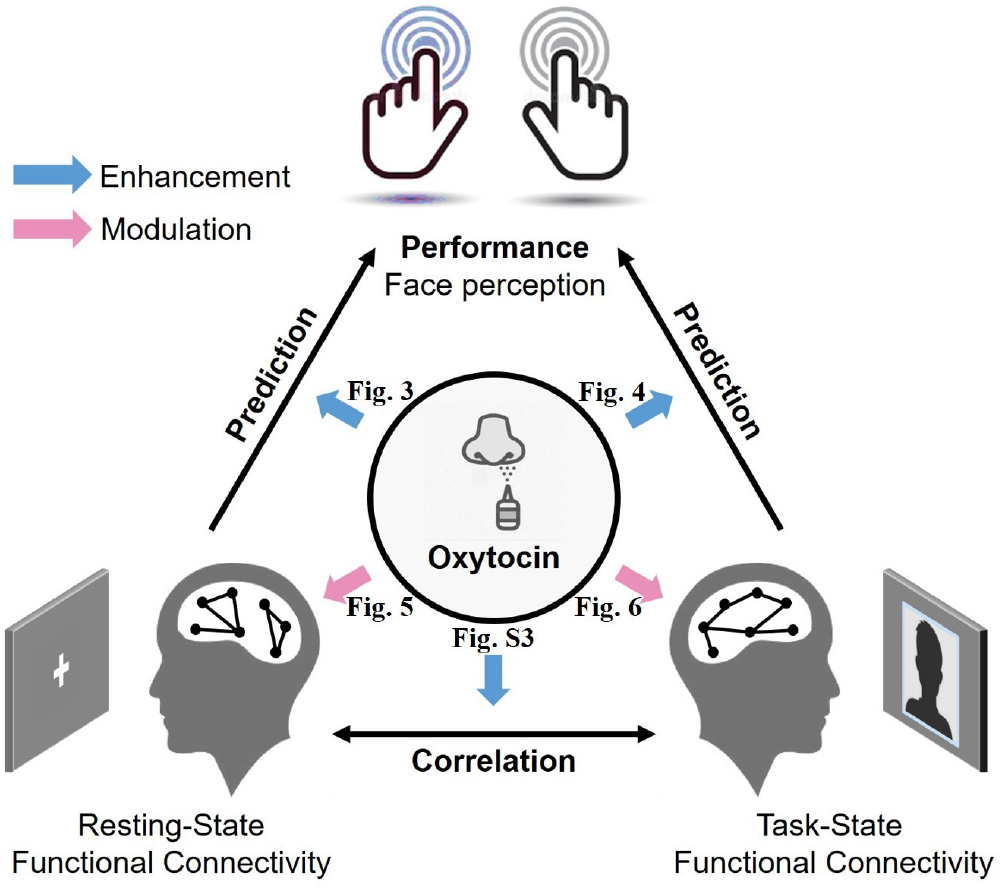
The framework of present study and corresponding result.

**Figure S2:**
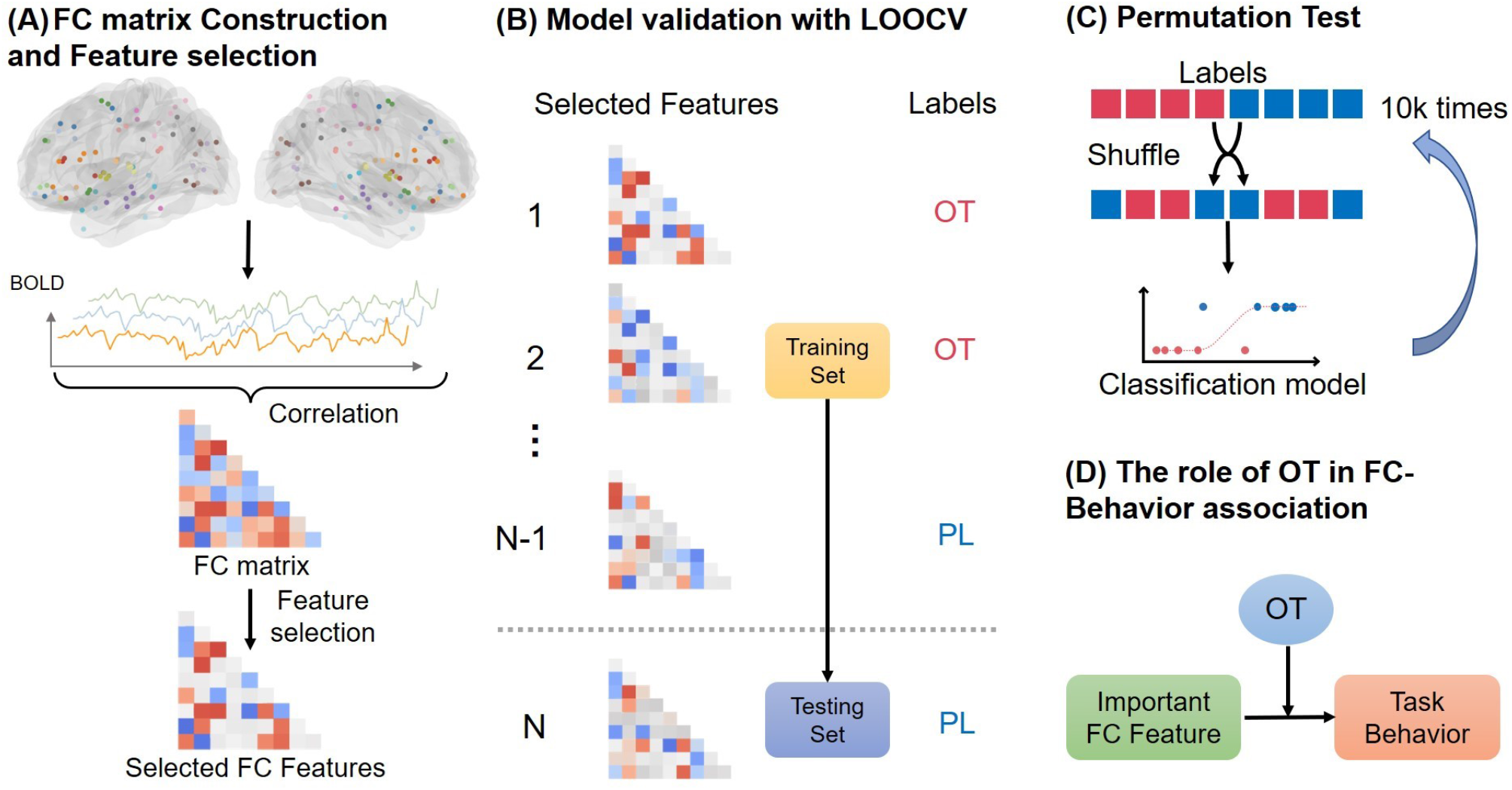
The workflow of the CPM predictor analysis for present study. (A) FC matrix construction and feature selection. (B) Model validation with LOOCV. (C) Test the performance of the predictor model. (D) Permutation test for statistics.

**Figure S3:**
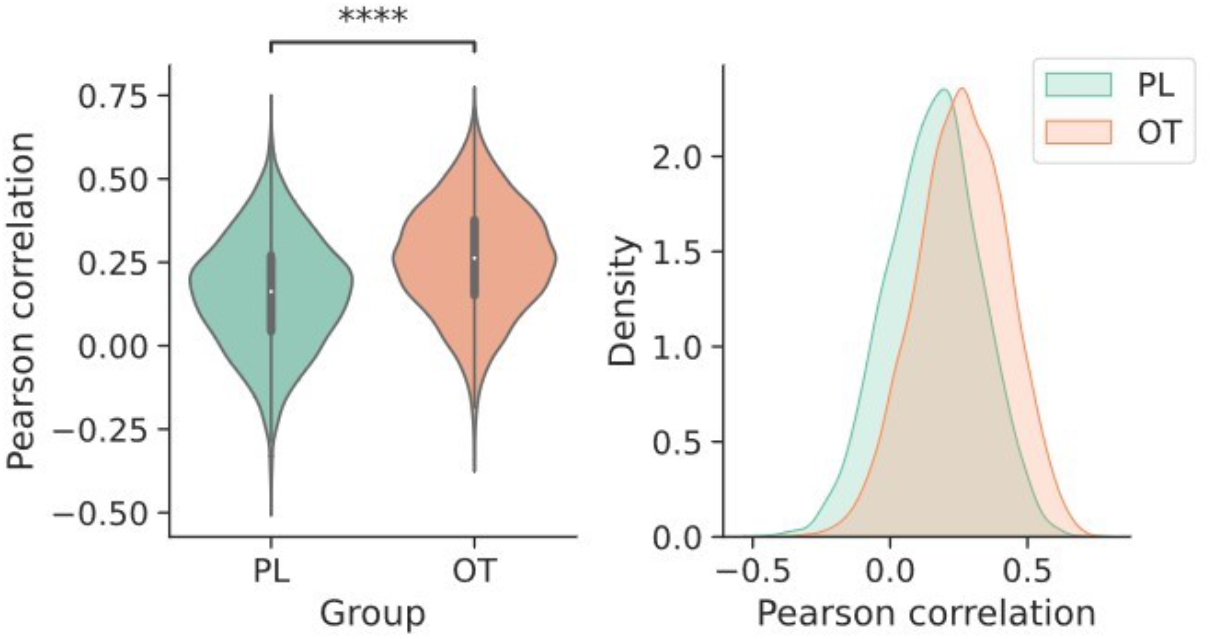
The Pearson correlation between each resting and task state functional connectivity edge for both groups. The correlation in OT group was significantly higher than PL group (*P <* 0.001).

**Figure S4:**
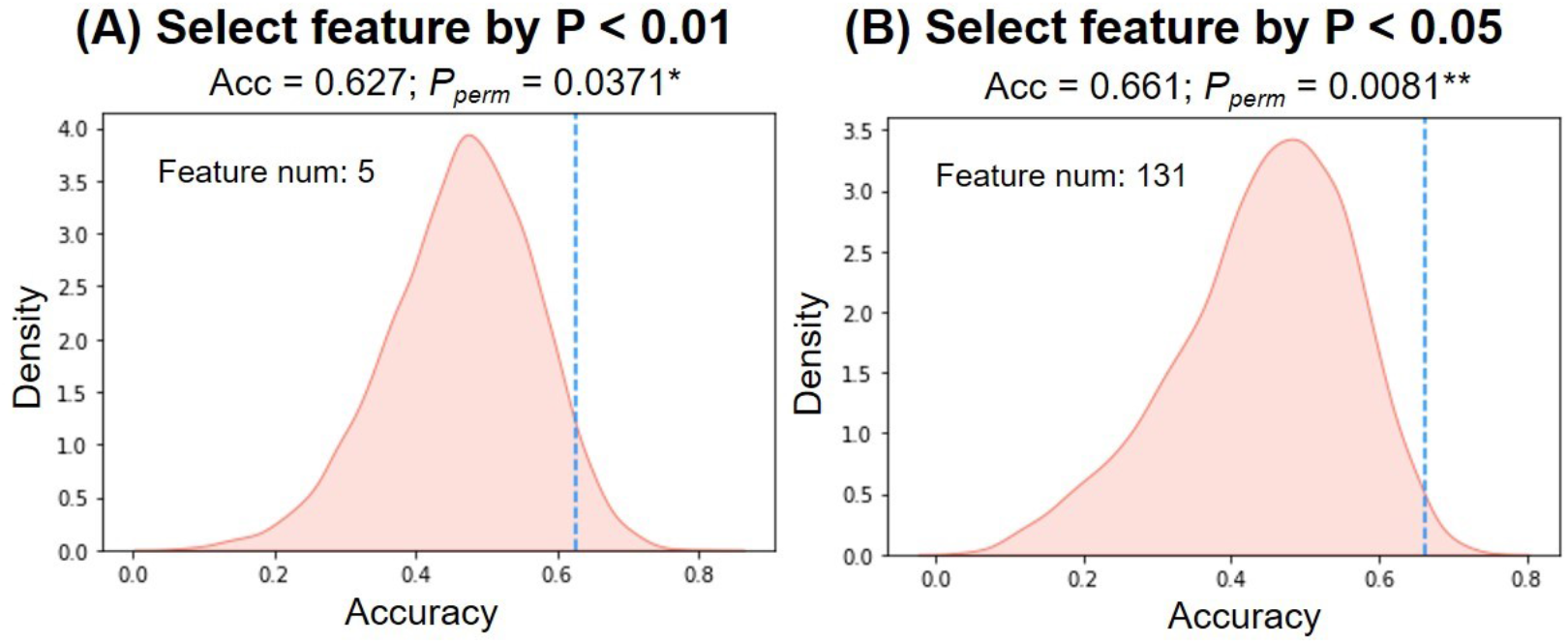
The result of rsFC classification using the different feature selected by different p-values. (A) 5 FC features during the feature selection with P *<* 0.01. (B) 131 FC features during the feature selection with P *<* 0.05.

**Figure S5:**
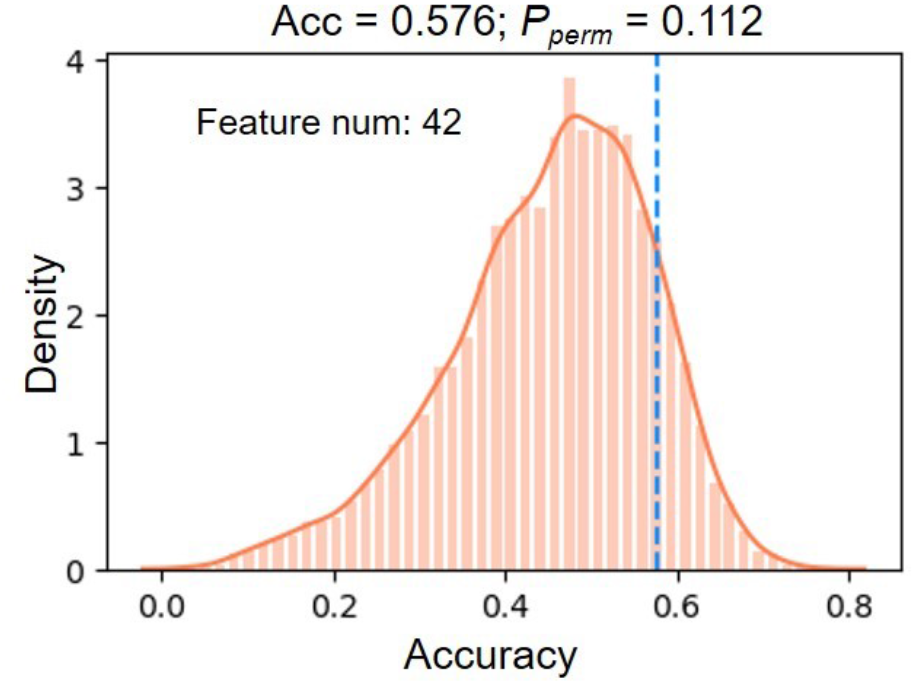
The result of tsFC classification using the different feature selected by different p-values = 0.05.

**Figure S6:**
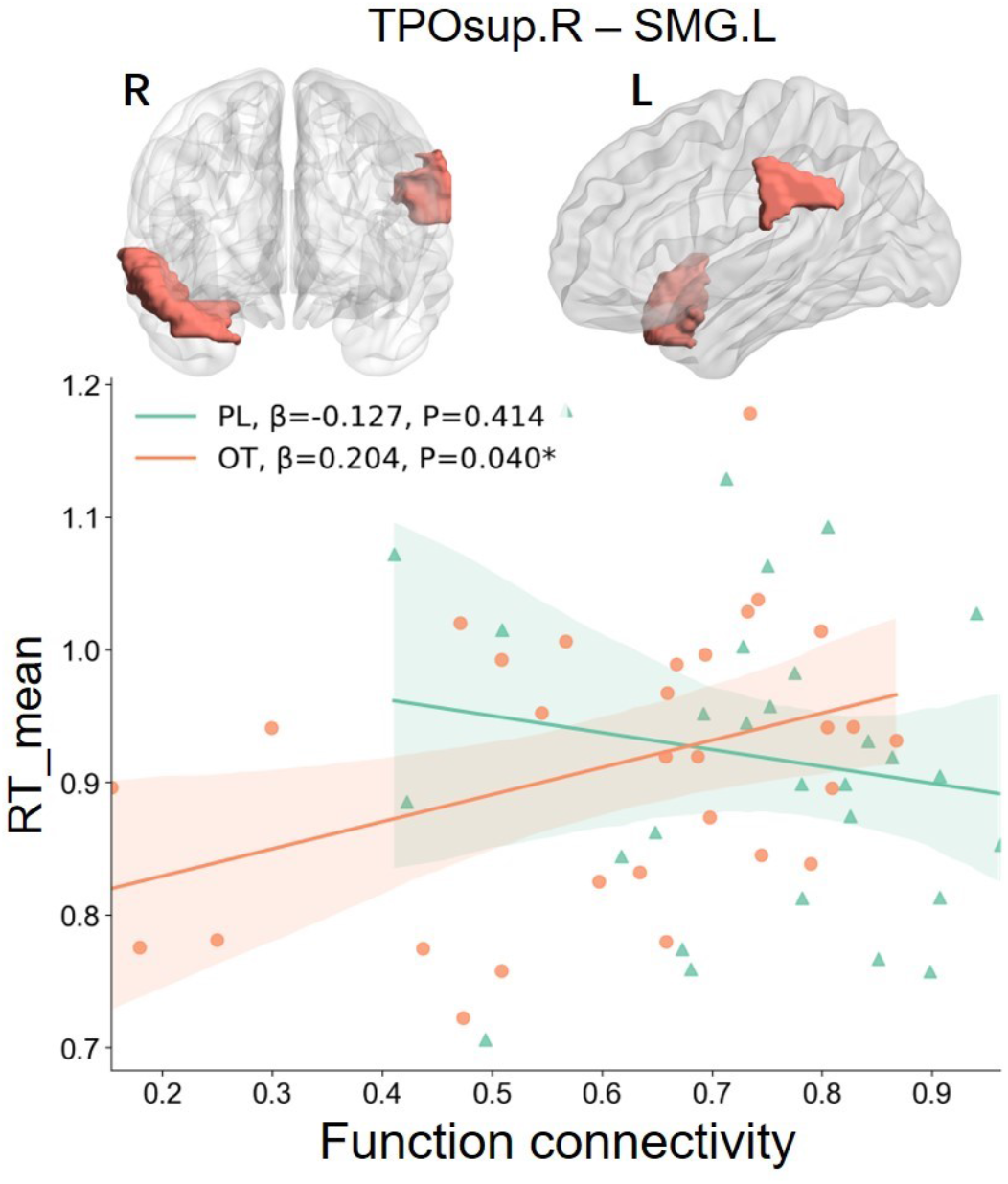
The result of SVM using the different feature selected by p-values = 0.05.

